# Assessing Age-Related Decline in Anti-Bacterial Immune Responses Using an Ex-vivo Assay in African Turquoise Killifish (*Nothobranchius furzeri*)

**DOI:** 10.1101/2023.11.24.568569

**Authors:** Gabriele Morabito, Francisco Daniel Dávila Aleman, Dario Riccardo Valenzano

## Abstract

Ex-vivo immune functional assays serve as essential tools for uncovering the intricate molecular mechanisms behind immune responses, both intrinsic and extrinsic, across a diverse spectrum of conditions and interventions. In this study, we devised an innovative assay aimed at quantifying anti-bacterial responses in immune cells from the primary immune organ of small teleosts. Our model of choice is the the African turquoise killifish (*Nothobranchius furzeri*), which has emerged as a prominent model system in the field of biology of aging, because of its natural short lifespan and the broad range of aging-related dysfunctions. Using our novel ex vivo assay, we tested age-associated differences in immune cell functionality, measured as anti-bacterial responses in young and aged turquoise killifish. Our results show that the ability to inhibit bacterial growth declines in cells extracted from aged killifish, compared to young killifish. This ex-vivo assay, designed for rapid measures of immune effector responses, holds the potential to be a scalable tool for assessing the cellular and molecular basis for anti-microbial immune responses under a range of interventions.

## Introduction

A key hallmark of immune health consists in the host’s ability to fend off pathogens and sustain a diverse commensal microbial community^1,2^. Conversely, aging is characterized by a declined capacity of the immune system to effectively fight pathogens and by significant changes in the microbiome, which could lead to “dysbiosis”^3,4^. Immune aging encompasses age-related dysfunction in immune organs, as well as alterations in the cellular dynamics of immune effector cells^5^. Immune aging has been suggested as a main systemic contributor to organismal aging^6^. Cell-type specific immune responses occurring during aging can be measured *in vitro*, e.g., assessing the age-dependent functional performance in phagocytosis, chemotaxis, antibody diversity or responses to proliferation signals ^7-12^. However, when cell-surface markers are not usable, such as in non-mammalian systems, immune cell functionality can be assessed from whole cell extracts derived from immune tissues (e.g. blood or hematopoietic niches). To study immune cells functionality during aging in non-mammalian animal models, we developed an ex-vivo assay that uses immune cells extracted from the primary hematopoietic organ (i.e. the kidney marrow) of the naturally short-lived African turquoise killifish (*Nothobranchius furzeri*)^13-15^. Turquoise killifish display several markers of spontaneous cellular immune aging at systemic, cellular and molecular level.

Recent findings showed that turquoise killifish display an age-related decreased diversity in the antibody repertoire, increased systemic inflammation, extensive markers of DNA damage and cellular senescence within progenitor cells of the primary immune niche^16,17^. However, compared to mammalian models, in turquoise killifish we still lack functional assays for immune cells. Unlike standard *in vitro* immune cells functional assays that test molecular features in specific cell types^18^ (such as phagocytosis in myeloid cells), we focused on developing an *in vitro* assay that measures a global response of freshly isolated primary immune cells, with the goal to capture a more holistic immune response against bacteria^19^. We adapted a previously published protocol that measures the anti-microbial properties of plasma isolated from the blood of various mammals^20^. In particular, we co-cultured bacteria with immune cells, rather than adding plasma, and we measured bacterial fluorescence, rather than optical density, over-time. In line with prior research demonstrating systemic and cellular markers of immune aging in turquoise killifish^11,17,21^, our *in vitro* bacterial growth assay underscores the age-dependent decline in the anti-microbial capabilities of immune cells extracted from the primary hematopoietic organ of these fish.

## Results

To quantify the ability of immune cells to inhibit bacteria growth, we developed an ex vivo functional assay (Bacterial Growth Inhibition Assay, BGIA), in which we co-culture fluorescent labelled bacteria with immune cells extracted from different immune organs. We aim to estimate the immune cell’s Bacterial Growth Inhibition Capacity (BGIC) by measuring bacterial growth in the presence and in the absence of immune cells (**Methods** and **Figure 1a-b**). First, we tested which killifish organ yields the highest number of surviving immune cells upon isolation. We found that cells isolated from the kidney marrow, the main hematopoietic niche of the adult killifish, resulted in high cell survival (80% of the total cells extracted) and successful cell isolation when compared with blood, gut and spleen (**Figure 1c**).

**Figure 1.**
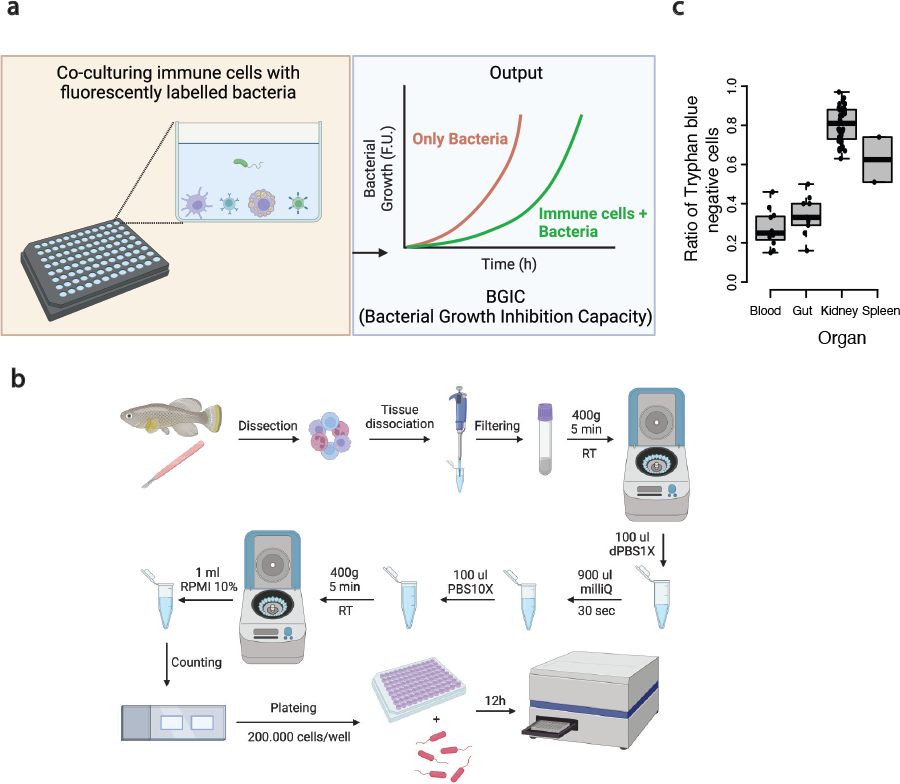
Bacterial Growth Inhibition Assay workflow. **a)** different immune cells cell-type specific functional features, inhibiting bacterial growth, are integrated together in a single index called BGIC by co-culturing immune cells with fluorescent-labelled bacteria. **b)** Schematics of the workflow used for immune cells isolation from the fish kidney marrows. **c)** Immune cell survival rate after the isolation procedure from different *killifish* organs.

Therefore, we selected kidney marrow as the main source of immune cells for the remainder of our study.

### Generation of fluorescently-labelled *Pseudomonas lundensis*

To rapidly and efficiently track bacterial growth over time, we genetically engineered bacteria to incorporate EYFP directly into the chromosome, ensuring long-term stability of the signal, as opposed to incorporating the construct in a plasmid. We used the bacterial Tn7 transposons system^22^, inserting the construct downstream to the “glmS” gene of *P. lundensis*, a bacterium we isolated from the turquoise killifish gut (**Figure 2a**). To test whether fluorescence intensity corresponded to bacterial growth, we compared bacterial fluorescence over-time with bacterial growth on plate. We performed a Colony Forming Unit (CFU) assay, a reliable measure of bacterial growth^23^, in parallel with measures of *P. lundensis* fluorescence. We measured bacterial growth in the presence of different concentrations of antibiotics that inhibit growth in gram negative bacteria ^24,25^ (i.e., kanamycin and streptomycin) (**Figure 2b-c**). Bacterial fluorescence over time significantly correlated with CFUs (**Figure 2d**), indicating that measured fluorescence can be used as a reliable measure of bacterial growth.

**Figure 2.**
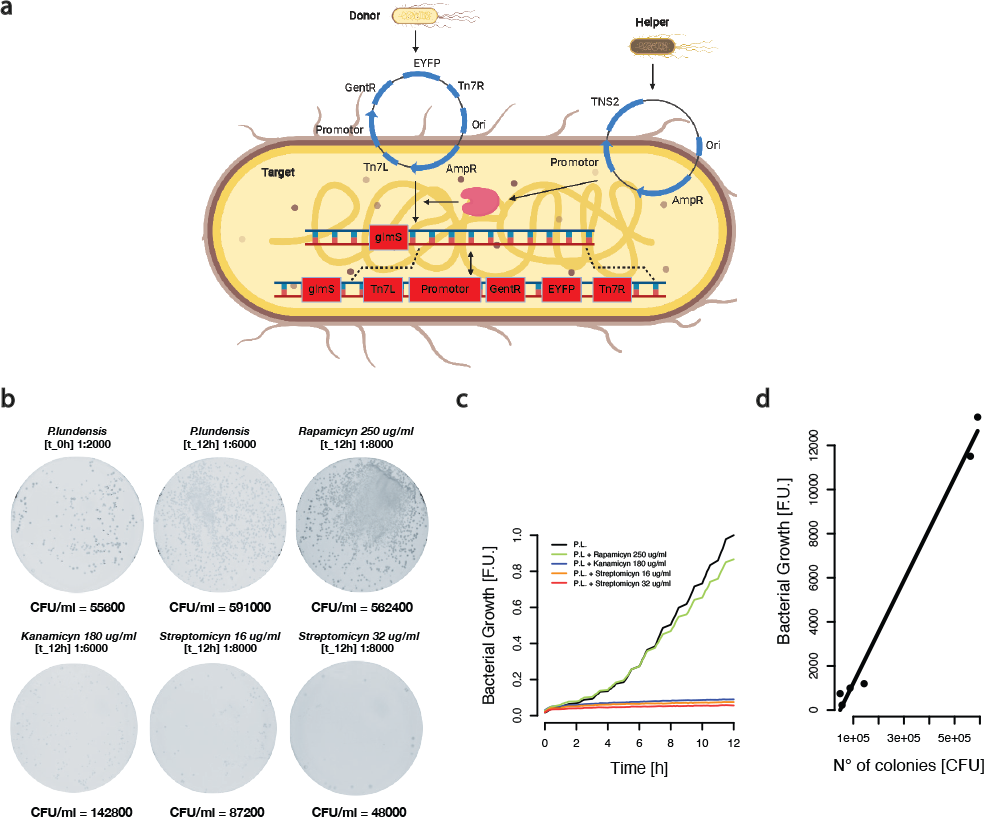
Tracking *Pseudomonas lundensis* cells divisions through EYFP fluorescence. **a)** Workflow used for the GentR / EYFP cassette stable insertion within *P*.*lundensis* chromosome. **b)** CFU assay using *P*.*lundensis* dilutions at 0 h and after 12 h incubation in presence of different antibiotics at different concentrations. **c)** *P. lundensis* growth curves, measured as fluorescence signal increase over-time, in presence of different antibiotics at different concentrations as was used in the CFU assay. **d)** Dotplot showing correlation between the number of colonies formed and the fluorescent signal obtained for each condition.

### Immune cells inhibit bacterial growth during BGIA

To test whether cultured immune cells can be activated after the isolation procedure, we ran an immune cell activation assay. Upon immune stimulation, activated immune cells form cell aggregates^26^. We isolated immune cells from the turquoise killifish kidney marrow and kept them for 24 h in presence or absence of 10 μg/ml peptidoglycan (PGN), a component of bacterial wall, which induces strong immune responses^27^. Our results showed the formation of immune cell aggregates only in the presence of PGN. These results are in line with previous observations in whole immune cells extracts after T cells activation^28^ and suggest that the cells extracted from the kidney marrow are immune cells that can be activated by bacterial antigens after the cell isolation procedure (**Figure 3a**).

**Figure 3.**
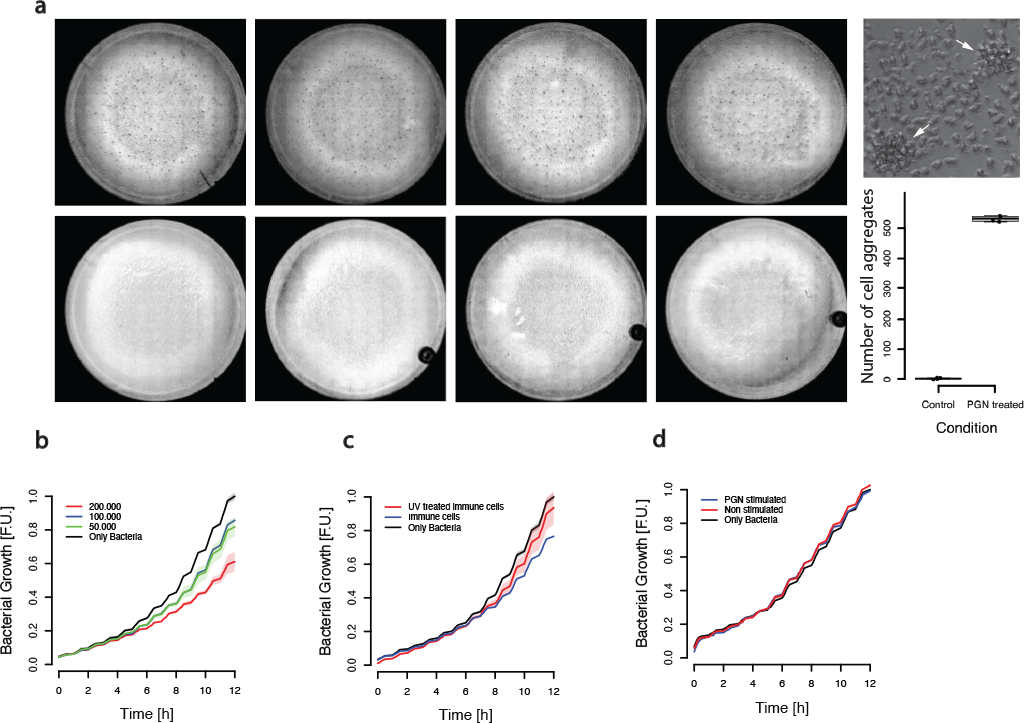
*Killifish* immune cells inhibit *P. lundensis* growth in vitro. **a)** Left: Bright field images of immune cells 24 h after isolation and incubation with or without 10ug/ml PGN. Each well contains 200.000 immune cells, 4 replicas per condition were used. Top right: bright field image of immune cell aggregates at 40X magnification. Bottom right: number of immune cell aggregates 24 h after the isolation procedure in wells incubated with or without 10ug/ml PGN. **b)** *P. lundensis* growth curves in presence of immune cells isolated from *killifish* kidney marrow and seeded in different amounts. **c)** *P. lundensis* growth curves in presence of 200.000 immune cells pre-treated with or without UV light for 30 min. **d)** *P. lundensis* growth curves in presence and absence of supernatant from overnight cultures of kidney marrows immune cells incubated or not with 10ug/ml PGN.

Next, we tested whether immune cells inhibit bacterial growth. First, we tested whether bacterial growth varies as a function of the number of the cultured immune cells (**Figure 3b**). We observed that *P. lundensis* growth was slower in the presence of higher numbers of immune cells.

We then asked whether external damage to immune cells impacted their effect on bacterial growth. We irradiated immune cells with UV-light for 30 min before performing the BGIA, which resulted in faster bacterial growth by irradiated immune cells, compared to the not irradiated control cells (**Figure 3c**). Overall, our results indicate that immune cells inhibit bacterial growth in culture and that damaged immune cells have a decreased capacity to inhibit bacterial growth.

To test whether soluble factors from the immune cell supernatant originating from the isolation procedure contributed to microbial growth inhibition, we cultured the extracted cells for 24 hours in the presence or absence of 10 ug/ml PGN, and then tested the impact of the resulting supernatants on *P. lundensis* growth (**Figure 3d**). Our results showed no impact on bacterial growth of cell supernatant from stimulated and un-stimulated immune cells, suggesting a negligible role of secreted factors on immune cell mediated inhibition of bacterial growth.

### An *Escherichia coli* strain for killifish ex-vivo immune cell testing

To test whether immune cells extracted from the killifish kidney marrow inhibit growth also in other bacteria than *Pseudomonas lundensis*, we used an *Escherichia coli* strain expressing chloramphenicol resistance (CmR) and GFP genes inserted within its chromosome (see **Methods**)^29^. Since we cultured killifish immune cells at 28 °C – corresponding to the temperature at which we grow killifish in our facility – we UV-mutagenized the CmR/GFP *E. coli* strain (**Methods** and **Figure 4a**) by selecting colonies growing at 28 °C with similar growing kinetics to *Pseudomonas lundensis* (**Figure 4b**).

**Figure 4.**
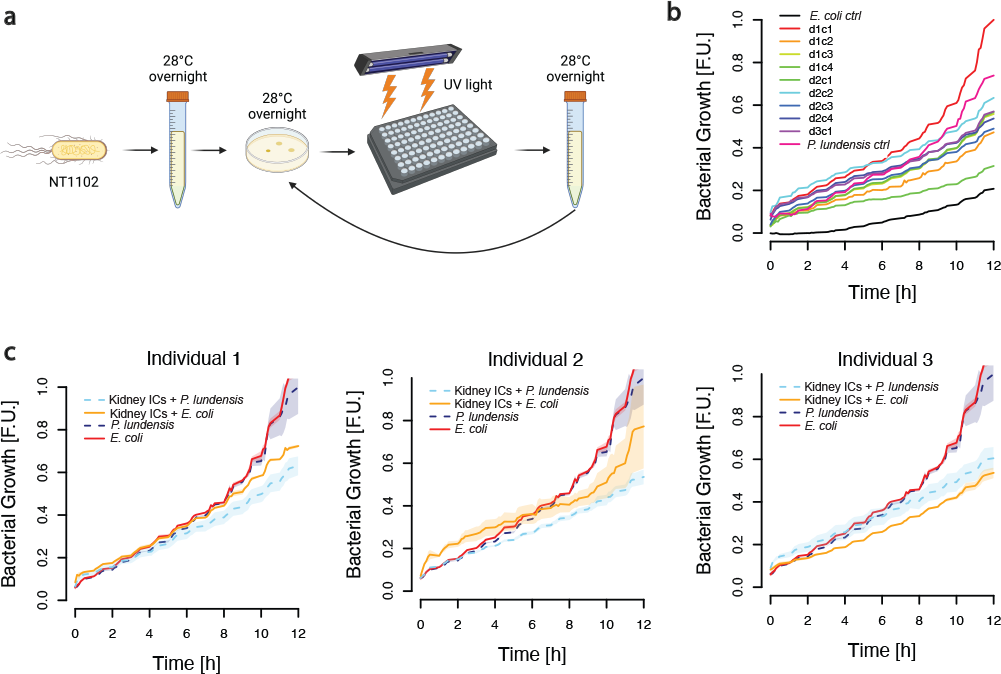
*Killifish* immune cells presence in the co-colture causes inhibition of *P*.*lundensis* growth: **a)** Workflow used for the UV random mutagenesis of fluorescent-labelled *E*.*coli*. **b)** Growth curves of several *E*.*coli* colonies (c1,c2,c3,c4) after different cycles of UV random mutagenesis (d1,d2,d3). **c)** *E*.*coli* and *P. lundensis* growth curves in presence of immune cells isolated from the kidney marrow of three different *killifish* of the same age.

### Growth inhibition rate of *P*.*lundensis* and *E*.*coli* by immune cells is individual dependent

To test if the growth of *P. lundensis* and *E. coli* is differentially inhibited in presence of immune cells, we tested the growth of both *P. lundensis* and *E. coli* in presence of immune cells of three different adult male killifish of the same age (13 week old). Immune cells from the three tested fish inhibited bacterial growth both in *P. lundensis* and *E. coli* (**Figure 4c**).

### BGIA reveal an age-dependent decrease of killifish immune cells capacity to inhibit bacterial growth

Since the extracted immune cells appear to inhibit bacterial growth *in vitro*, we next asked whether immune cells extracted from killifish at different ages had different impacts on bacterial growth, compatibly with results showing that killifish aging has several features of immune aging^11,17^. We ran the assay using labelled *P. lundensis* and fixed immune cell numbers (200.000 per well) isolated from the kidney marrows of four 7 week old (young-adult) and six 16 week old (aged) killifish (**Figure 5a**).

**Figure 5.**
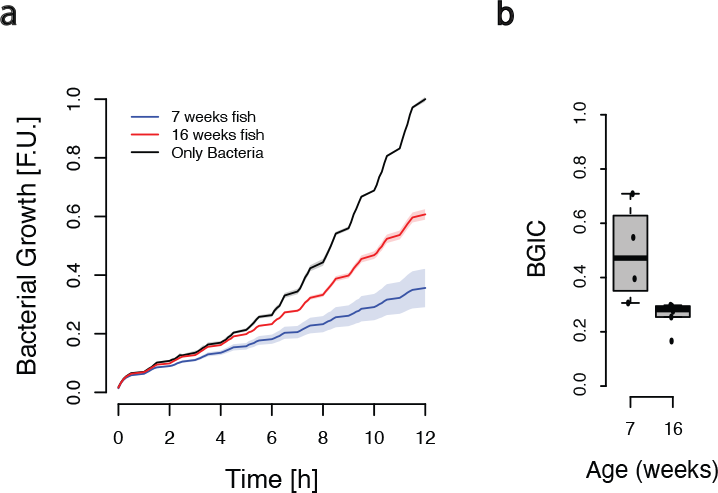
*Pseudomonas lundensis* growth inhibition by young and aged immune cells isolated from *killifish* hematopoietic niche: **a)** On the left growth curves of *P*.*lundensis* in the presence of none, young (7 week old) or aged (16 week) *killifish* immune cells. On the right boxplots showing BGIC quantification from the previous curves at the two age timepoints. **b)** Boxplots showing BGIC age-dependent decline in immune cells isolated from hematopoietic niches of *killifish* at different ages: young (7 week old), adult (11 week old), aged (16 week old) and geriatric (19 week old). BGIC values measured in the same experimental run are indicated by the same colour of the dots. All the statistics were done using non-parametric Wilcoxon ranked sum test.

Immune cells from the young killifish cohort inhibited bacterial (*P. lundensis*) growth more than immune cells from the old killifish cohort. To further test consistency and repeatability of the age-dependent decline in the bacterial growth inhibition capacity of killifish immune cells, we compared the BGIC obtained in different runs in which we used immune cells isolated from the kidney marrows of *killifish* at different ages (**Figure 5b**). We observed a consistent decrease in the BGIC as a function of the age of the immune cell donor.

## Discussion

Functional assays that test the functionality of immune cells are designed to evaluate the capacity of specific immune cells types to perform specific functions critical to the immune response. The importance of functional assays in immunology lies in their ability to provide molecular insights into immune system’s performance, which is essential for understanding immunological processes, disease mechanisms, and therapeutic development. However, conventional functional assays often rely on antibodies that facilitate sorting of specific immune cell types, becoming un-accessible to model systems that lack available commercial antibodies, e.g., non-mammalian systems. Our work addresses the need for alternative functional assays that can be applied to non-mammalian systems, such as teleosts.

Here we developed a novel functional *in vitro* assay (BGIA) that quantifies complex immune responses in immune cell types obtained from crude immune cells extracts of the turquoise killifish kidney marrow, which contains all the major vertebrate immune cell types ^17^. Our assay measures bacterial growth in the presence of extracted immune cells, used as a proxy of immune cell function. Testing bacterial growth in the context of young and old-derived *killifish* immune cells shows an age-dependent declined capacity to inhibit bacterial growth. These results are in line with previous work showing that turquoise killifish undergo spontaneous and extensive aging of the immune system ^11,17,21^. Together, BGIA represents a promising functional assays to rapidly test immune responses *ex vivo* in non-canonical model organisms.

## Material and Methods

### Animal housing and euthanasia

Turquoise killifish (GRZ-AD strain) were maintained in plastic fish tanks within a recirculating aquaculture system, adhering to the specifications outlined in holding license 11-003798 at the Leibniz Institute on Aging, Fritz-Lipmann Institute. Euthanasia of the fish was performed by immersion in a 1.5 g/L Tricaine solution, followed by cutting at the caudal peduncle using a scalpel.

### Blood Collection

To extract blood, the euthanized fish were placed with the amputated caudal portion immersed in a 50 μl drop of Blood Collection Buffer (dPBS1X + 1:500 diluted solution containing Gentamicin 5 mg/ml and Amphotericin B 125 μg/ml) within a Petri dish. Subsequently, blood was collected into a 1.5 ml Eppendorf tube.

### Extraction of Immune Cells

To extract immune cells, kidneys, gut and spleen were collected in a 1.5 ml Eppendorf tube containing 900 μl of 1X DPBS. The gut underwent pre-treatment by incubating it for 20 minutes at 37 °C in 1 ml of Collagen Digestion Buffer (1X DPBS + Collagenase 0.75 mg/ml + heat-inactivated FBS 5%). Tissues were homogenized by gently pipetting into the Eppendorf tube. To isolate immune cells from epithelial cells, the sample solution was filtered through a 35 μm nylon membrane in the cap of a 5 ml FACS tube, followed by immediate transfer into a new 1.5 ml Eppendorf tube according to a previously established protocol^30^. Next, to enrich the samples for immune cells over red blood cells, the immune cells suspensions from kidney, spleen and gut were centrifuged at 400g and room temperature for 5 minutes. The supernatant was discarded, and the pellets were resuspended in 100 μl of 1X DPBS (blood sample was already resuspended in 100ul of 1XDPBS from blood collection). Afterwards, to cause an osmotic stress that allow only immune cells in the sample to survive, 900 μl of milliQ water was added, samples were immediately gently inverted for 20 seconds, followed by the immediate addition of 100 μl of PBS10X. This mixture was then centrifuged at 400 g and room temperature for 5 minutes. Finally, the supernatant was discarded, and the immune cell pellet was resuspended in RPMI 10% medium (RPMI + Hepes 1:40 + heat-inactivated FBS 10% + 1:500 diluted solution containing Gentamicin 5 mg/ml and Amphotericin B 125 μg/ml).

### Cell concentration and viability measurement

To assess immune cells survival after the extraction, a 10 μl aliquot of Trypan Blue and 10 μl of the cell suspension sample were combined in a 1.5 ml Eppendorf tube. The resulting solution was gently pipetted, and 10 μl of this mixture were loaded into a well on the nCounter slide (Bio-rad, ref: 1450011). Subsequently, the nCounter slide was inserted into the TC20 Automated Cell Counter (Biorad TC20TM, ref: 1450102) to assess the cell viability and concentration of the sample.

### Fluorescent labeling of P. lundensis

Single colonies of *E. coli* SM10 pEYFP:GntR, *E. coli* SM10 pTNS2:AmpR, and *P. lundensis* were selected from solid cultures. Each colony was added to separate 10 ml Falcon tubes containing 5 ml of LB broth, with specific antibiotic solutions based on the strain. Sub-cultures were initiated, and upon reaching the exponential phase (OD600 of ∼0.4-0.6), bacterial cultures were placed on ice. To transform *P. lundensis* through bacteria conjugation, the bacteria subculture was co-cultured in a mixed bacterial suspension with the two *E* .*coli* subcultures within a 1.5 ml Eppendorf tube. Then, the sample was centrifuged at 7000g room temperature for 2 min. and the pellet was resuspended in 25 μl of Washing Solution (H2O milliQ + NaCl 0.7%), and subsequent filtration onto a 0.45μm filter disc. Next, the trapped bacteria were eluted in 1 ml of Washing Solution and the resulting cell suspension was collected^31^. To isolate the labeled *P. lundensis* the collected suspension was incubated in LB broth with Gentamicin 5 mg/ml and Amphotericin B 125 μg/ml inside overnight.

### UV Random Mutagenesis on E. coli

To generated a labeled *E. coli* strain that show an optimal growth at 28 °C we incubated overnight *E. coli* NT1102 strain, transformed with pJLRCS-GFP plasmid and obtained by Prof. Georg A. Sprenger at University of Stuttgart (Germany), in LB broth containing Chloramphenicol 25 μg/ml. After centrifugation and resuspension, a portion was seeded on an agar plate with Chloramphenicol and incubated overnight at 28 °C. The colony with the best growth was selected, and UV mutagenesis was performed for 30 min. The mutagenized bacteria were subjected to 4 total cycles of selection and incubation, and the selected colonies were assessed for GFP fluorescence using a plate reader.

### Bacterial Growth Assay

Engineered *P. lundensis* (GentR:EYFP) was cultured overnight in LB broth with specific antibiotics. On the day of the assay, immune cells were freshly isolated, and approximately 2x10^5^ cells/well were added to a 96-well plate with RPMI 10% medium. The bacterial culture was centrifuged, and the OD_600_-adjusted (in the range of 2-2.3) bacterial resuspension was added to the wells. The plate was then placed in a plate reader to measure fluorescence intensity in the EYFP/GFP channel every 10 minutes for 12 hours, with shaking (300rpm) before each measurement cycle. Negative and positive controls were included for each immune cell biological replica.

## Acknowledgments

We are thankful to all members of the Valenzano lab at the Max Planck Institute for Biology of Ageing and at the Fritz Lipmann Institute (FLI), Leibniz Institute on Aging, for the support and continuous input on this project. We thank the fish facility staff, both at the Max Planck Institute for Biology of Ageing, as well at the FLI, for their continuous support of our research. We thank Professor Georg A. Sprenger and his team at the University of Stuttgart for providing us with a GFP-labelled *E. coli* strain. This project was supported by the Max Planck Society, by the Leibniz Association, and by the DFG grant CRC1310.

